# The effect of stride length on lower extremity joint kinetics at various gait speeds

**DOI:** 10.1101/363788

**Authors:** Robert L. McGrath, Melissa L. Ziegler, Margaret Pires-Fernandes, Brian A. Knarr, Jill S. Higginson, Fabrizio Sergi

**Affiliations:** Department of Biomedical Engineering, University of Delaware, Newark, DE 19713, USA; Biostatistics Core, College of Health Sciences, University of Delaware, Newark, DE 19713, USA; Department of Biomedical Engineering, University of Florida, Gainesville, FL 32611, USA; Department of Biomechanics, University of Nebraska, Omaha, NE 68182, USA; Department of Mechanical Engineering, University of Delaware, Newark, DE 19713, USA

## Abstract

Robot-assisted training is a promising tool under development for improving walking function based on repetitive goal-oriented task practice. The challenges in developing the controllers for gait training devices that promote desired changes in gait is complicated by the limited understanding of the human response to robotic input. A possible method of controller formulation can be based on the principle of bio-inspiration, where a robot is controlled to apply the change in joint moment applied by human subjects when they achieve a gait feature of interest. However, it is currently unclear how lower extremity joint moments are modulated by even basic gait spatio-temporal parameters.

In this study, we investigated how sagittal plane joint moments are affected by a factorial modulation of two important gait parameters: gait speed and stride length. We present the findings obtained from 20 healthy control subjects walking at various treadmill-imposed speeds and instructed to modulate stride length utilizing real-time visual feedback. Implementing a continuum analysis of inverse-dynamics derived joint moment profiles, we extracted the effects of gait speed and stride length on joint moment throughout the gait cycle. Moreover, we utilized a torque pulse approximation analysis to determine the timing and amplitude of torque pulses that approximate the difference in joint moment profiles between stride length conditions, at all gait speed conditions.

Our results show that gait speed has a significant effect on the moment profiles in all joints considered, while stride length has more localized effects, with the main effect observed on the knee moment during stance, and smaller effects observed for the hip joint moment during swing and ankle moment during the loading response. Moreover, our study demonstrated that trailing limb angle, a parameter of interest in programs targeting propulsion at push-off, was significantly correlated with stride length. As such, our study has generated assistance strategies based on pulses of torque suitable for implementation via a wearable exoskeleton with the objective of modulating stride length, and other correlated variables such as trailing limb angle.

## Introduction

Robot-assisted training is a promising tool under development for improving walking function [1, 2]. A primary indicator of gait performance improvement is gait speed (GS), which is associated with a better quality of life [3] and overall functional status [4]. Currently, it is not well understood how the modulation of assistance provided by a robot during gait training will lead to changes in GS. The gait parameter of GS is known to be correlated with anterior-posterior ground reaction force, the propulsive force of the foot against the ground. Furthermore, propulsive impulse, the propulsive force integrated over time, is associated with the posture of the trailing limb at push-off [5]. The posture of the trailing limb at push-off is quantified by one kinematic parameter, known as trailing limb angle (TLA), defined as the angle of the line connecting the hip joint center and foot center of pressure at the instant of peak propulsive force, relative to the global vertical axis [6]. In healthy control subjects, it was observed that when increasing GS, the increase in TLA contributes twice as much as the increase in ankle moment to the resulting increase in propulsive force [7]. In older adults exposed to a biofeedback paradigm for increasing propulsive force, increased in TLA and decreased hip flexor power were the primary means by which they increased propulsive force [8]. Therefore, TLA has been advanced as a variable of interest for robot-assisted gait training paradigms aimed at improving walking function.

Our research group is exploring the use of robot-assisted gait training to directly target and modulate TLA, thus allowing subjects to modulate GS. A possible controller could be composed of torque pulses applied at specific instants during the gait cycle, with the advantage of not constraining gait to follow prescribed trajectories [9]. This approach has been shown in previous studies to be a successful method of robot-assisted gait training [10, 11]. However, in absence of models of the human response to a robotic input, it would be difficult to define parameters for such a controller acting on multiple degrees of freedom. A possible method of controller formulation can derive from the principle of bio-inspiration, where a robot is controlled to apply the difference in joint moment applied by human subjects when they achieve a desired gait feature (in this case modulation of TLA), relative to their normal walking condition. Once the effects of the variable of interest have been identified, a rehabilitation robot could be controlled in either assistive, resistive, or perturbation mode to deliver different forms of robot-assisted training [12]. To support the development of such a controller, we first required knowledge about the joint moments applied by healthy control subjects to modulate TLA at a range of GSs. Since TLA has not been a primary measure of interest in the biomechanics literature, we extended our search to a more common variable, likely correlated to TLA, such as stride length (SL).

The joint kinetics associated with GS modulation in healthy control subjects have been thoroughly elucidated in the literature, where an increase in GS is generally associated with increase in magnitude of peak joint moments. A very early investigation examining knee kinetics found increasing GS to be strongly correlated with an increase in peak knee extension moment [13]. Further work found peak hip flexion and extension moments, knee flexion and extensions moments, and ankle plantar and dorsiflexion moments to all increase with GS. However, these changes in joint kinetics to increasing speed were primarily observed at the hip, particularly in extension, and secondarily at the ankle for purposes of support [14]. Recently, an increase in GS was observed to be associated with an increase in peak hip extension moment during loading response, knee flexion moment in late stance and peak ankle plantarflexion moment [15]. Also, hip extension during loading response, hip flexion during early swing, hip extension during late swing, knee flexion during loading response, knee extension during early stance and ankle dorsiflexion during loading response all increased with increases of gait speed and stride length [16].

However, fewer investigations have focused on the joint kinetics associated with the modulation of spatiotemporal parameters such as SL or TLA, and studied this effect at multiple GS values. Summed joint work has been observed to be strongly correlated with SL in both young and old adults, where young adults primarily utilized swing phase hip work to modulate SL and old adults utilized ankle and knee joint work [17]. An early investigation found stance phase peak knee extension moment and peak knee flexion moment to be strongly correlated with increasing SL [13]. A more recent and in-depth investigation found that as SL increased, peak ankle plantarflexion moment, plantarflexion moment at 40% of stance, and peak knee extension moment all increased, while peak knee flexion moment and peak hip flexion moment decreased [18].

Thus far, the factorial modulation of both GS and SL and resulting hip, knee, and ankle kinetics has not been investigated; as such it is unclear how lower extremity joint moments are modulated by both gait parameters. Addressing this gap of knowledge, we designed an experimental study to establish the effects of GS and SL on the resulting hip, knee, and ankle joint moments. The findings are intended to inform the design of a robotic controller that delivers pulses of torque to the lower extremity joints with optimal timing and amplitude to induce desirable modulations of GS and SL, and of associated spatiotemporal parameters.

## Materials and methods

### Subjects

20 healthy adults (10 males, 10 females) were recruited to participate in this study (protocol approved by the University of Delaware Institutional Review Board, protocol no. 619724). Subjects — age (mean ± std) 21.55 ± 2.50 yrs, height 1.73 ± 0.08 m, body mass 69.20 ± 8.73 kg — were naive to the purpose of the study, and free of orthopedic or neurological disorders affecting walking function. Subjects were required to wear their own comfortable athletic shoes and lightweight clothing for the walking experiment.

### Setup

Subjects walked on an instrumented dual-belt treadmill (Bertec Corp., Columbus OH, USA), as shown in Fig. 1,while wearing thirty-six reflective spherical markers (4 on the pelvis, 4 per thigh, 4 per shank, 2 per knee, and 6 on ankle/foot). An eight camera Raptor-4 passive motion capture system (Motion Analysis Corp., Santa Rosa CA, USA), for subjects 1-14, and ten camera Vicon T40-S passive motion capture system (Oxford Metrics, Oxford, UK), for subjects 15-20, were used to measure marker position in space. Marker data were acquired at 100 Hz, while the treadmill analog force/torque data were acquired at 2 kHz. A 24-in screen was placed at approximately 1500 mm anteriorly from the center of the treadmill, and was used in biofeedback conditions. The screen provided visual feedback about the SL measured at the previous gait cycle (starting and ending at right heel strike) which was updated within 20 ms after each right heel strike. In this experiment, SL for cycle *k* was defined based on the right heel strike time *t* and anteroposterior coordinate in the laboratory frame *x* and constant velocity of the treadmill *v*:

**Fig 1.**
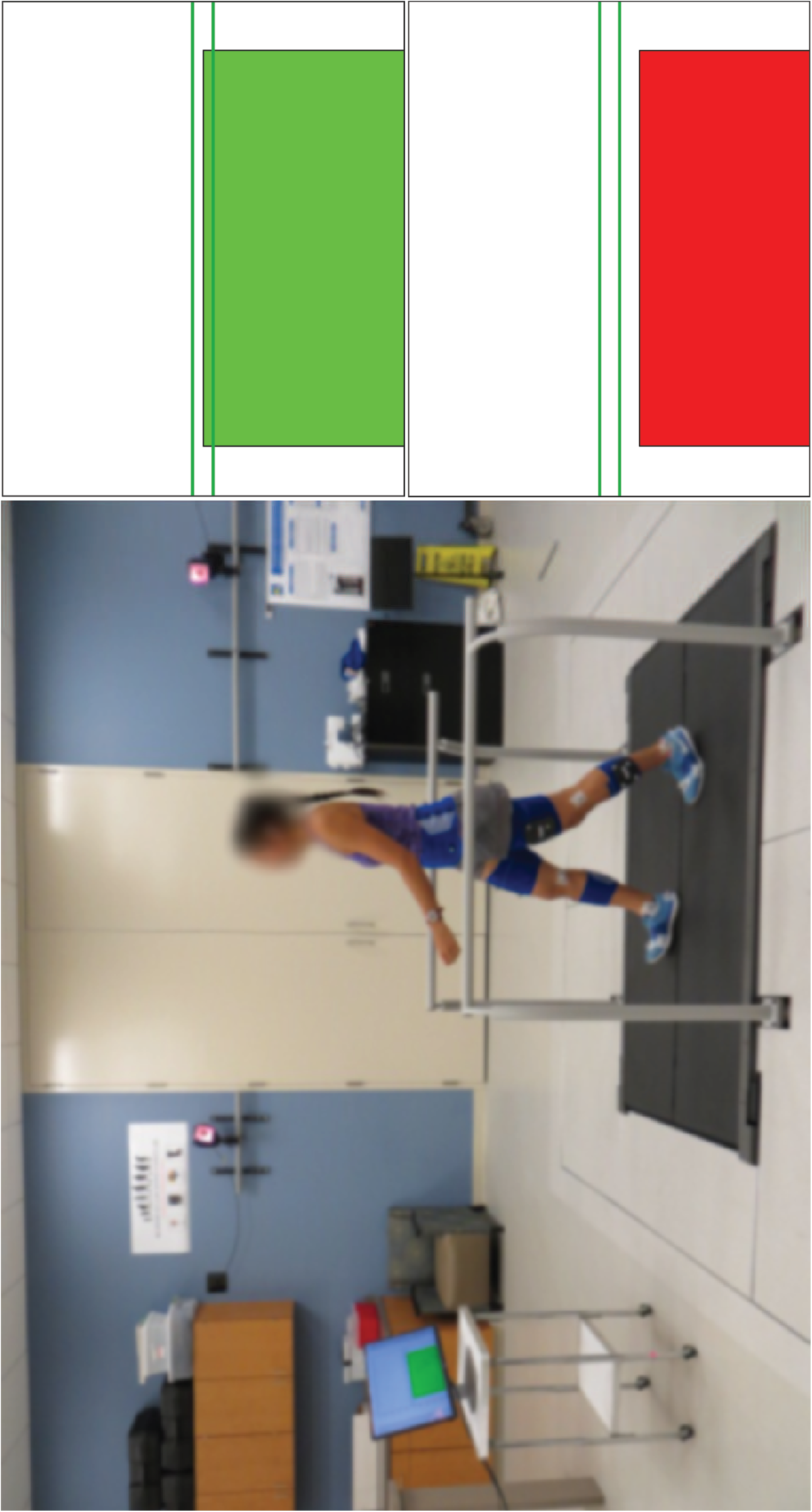
The experimental setup - subject walking on instrumented treadmill while wearing retroflective markers captured by infrared camera system. Real-time visual feedback provided on screen bar height and color cues to impose stride length condition. A green bar indicated a stride length within target range (horizontal lines) and a red bar indicated stride length outside of target range.

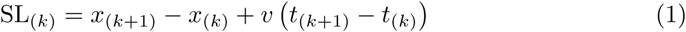

Visual feedback of SL_(*k*)_ was provided in terms of the height of a bar, while the desired SL was displayed as a horizontal line with dashed lines indicating the ± 10% range. The bar indicating SL_(*k*)_ was color coded to indicate whether the measured value was within ± 10% of the desired value. During biofeedback sessions, subjects were instructed to modify the length of their strides to achieve the target range, while walking at treadmill-imposed speeds.

### Procedures

Subjects were exposed to a total of fifteen experimental conditions, determined as the combinations of two factors: *i)* GS, with five levels, and *ii)* SL, with three levels. Factor levels were defined in terms of percent change relative to subject self-selected (ss) values to accommodate inter-subject variability in gait parameters. Moreover, to account for the correlation between GS and SL [19], we first measured self-selected stride length (ss-SL) at all speeds, and defined biofeedback-modulated SL conditions as percent changes of SL relative to the ss-SL at any given speed. This experimental setup allowed us to investigate joint kinetics underscoring an increase or decrease of SL relative to the subject’s self-selected stride length, at all speeds.

### Self-selected gait speed

A preliminary set of trials were conducted to calculate the subject’s self-selected gait speed (ss-GS). Subjects were asked to walk on the treadmill moving at an initial speed of 0.5 m/s, with the treadmill speed gradually increased by intervals of 0.03 m/s, and to indicate when ss-GS was reached. The same procedure was repeated by starting with the treadmill at 1.8 m/s, and decreasing treadmill speed in increments of 0.03 m/s, until the subject indicated that ss-GS had been reached. This procedure was repeated three times and the ss-GS was calculated as the average between the six measured treadmill speed values.

### Non-biofeedback conditions

After determination of ss-GS, five walking trials were conducted consisting of ninety seconds of acquired data in the absence of biofeedback. In each trial, treadmill speed was imposed at one of five percentages of the subject’s ss-GS [80%, 90%, 100%, 110%, 120%] in a randomized order. For each GS, ss-SL was calculated as the mean SL measured at that treadmill speed and utilized for the definition of subsequent desired SL values at each GS.

### Biofeedback conditions

After determination of ss-SL for all five GS conditions, ten additional walking trials were conducted consisting of ninety seconds of data acquisition, two for each treadmill speed value, using biofeedback to cue a desired SL. For each GS, the desired SL was set to be either 17% greater or 17% smaller than the ss-SL at that GS, in a random order. The range of change in SL values was specified based on previous studies showing feasibility of achieving distinguishable gait kinetics when SL was modulated by 17% of the ss value [19]. The investigator initiated data acquisition for each condition when the subject sufficiently achieved the cued SL condition specified via biofeedback.

## Data analysis

### Processing

Raw marker trajectories were labeled offline. Marker position and force/torque data were fed into a standard Visual3D processing pipeline, which included *i)* noise gating of measured force with a 25 N threshold *ii)* low-pass filtering of marker and force/torque data (Butterworth filter at 6 Hz and 30 Hz cut-off frequency, respectively), *iii)* interpolation of missing marker data with a third order polynomial fit for a maximum gap size of five samples, *iv)* application of the subject mass, height, and standing calibration marker positions to an anatomical template to derive a 7 segment (pelvis, thighs, shanks, and feet) link-segment subject model, and *v)* use of an optimization algorithm applied to inverse kinetics and kinematics equations to obtain joint angles and net moments.

In a custom MATLAB (MathWorks, Natick, MA, USA) script, hip, knee, and ankle joint angles and moments for the right leg in the sagittal plane were extracted and filtered with a 2^*nd*^ order low-pass zero-shift Butterworth filter with a cut-off frequency of 15 Hz. Gait cycles were segmented between subsequent heel strike events, defined as the instants at which the vertical ground reaction force changed in value from zero to positive, and remained positive for a minimum of 400ms. Due to events such as marker occlusion or subjects’ foot stepping on the contralateral force plate, acquired data were manually screened and some gait cycles were excluded from the analysis. A minimum of 25 segmented gait cycles were linearly resampled in the [0,100] % gait cycle domain 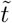 and averaged at each point in gait cycle to yield an average hip, knee, and ankle joint moment profile 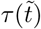 for each of the 15 experimental conditions for each of the 20 subjects.

Prior to obtaining average group moment profiles, joint moment profiles were non-dimensionalized. In agreement with [20], the non-dimensional joint moment 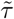 was calculated for each joint as 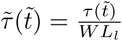, where *W* is body weight in N, and *L*_*l*_ is leg length in m, measured as the distance between the hip joint center and the floor during straight-leg standing.

### Protocol validation

A non-dimensional GS was defined as the Froude number 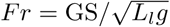, where *g* is the acceleration of gravity. Although several other factors such as body mass and athletic fitness condition account for the variability in ss-GS across individuals [21], the Froude number has been extensively used to describe the conditions underlying the transition from walking to running in several species [22]. As such, we used the Froude number as an index of across-subject dynamic similarity in ss-GS: a smaller variance of Froude numbers within a group of individuals should reflect consistent gait kinetics. We calculated the coefficient of variation 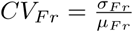 as the ratio between the standard deviation and the mean of Froude numbers corresponding to the ss-GS condition, and compared it to alternative indices, 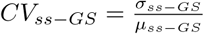 that uses ss-GS, and 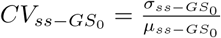 that uses ss-GS normalized by leg length.

Two gait parameters were also calculated; SL was measured using eq. (1), while TLA was calculated as the angle relative to the vertical axis of the line connecting the hip joint center and the position of the center of pressure at the instant of maximum anterior ground reaction force. To determine if the imposed biofeedback effectively modulated SL in healthy subjects, we assessed the distribution of SL values across all three feedback conditions at the five different gait speeds. We calculated the mean TLA and SL_0_ value for each of the 15 conditions for all subjects and conducted a linear correlation analysis using these two measurements as factors to assess the statistical association between normalized stride length and TLA.

### Continuum analysis

We sought to determine if the two factors GS and SL had any significant effect on the sagittal plane moment profiles for the hip, knee, and ankle joint 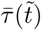, and, if so, at which phase of a gait cycle was a significant effect of either factor measured. We conducted an analysis for the main effects of the two factors, GS and SL, by implementing a repeated-measure 2-way ANOVA on the mean joint moment profiles measured from each subject and experimental conditions, spanning exhaustively the 15 combinations of factors. ANOVA was conducted to test the null hypothesis that neither factor (GS and SL), nor their interaction, induce a significant effect on joint moment at any time point.

Since the dependent variable 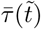 is one-dimensional (1D) smoothed time series including highly temporally correlated data, and not a zero-dimensional scalar quantity (e.g. peak torque, range of motion, etc.), definition of confidence intervals and control of false positive rates (FPR) requires proper correction for multiple comparisons that accounts for the temporal correlations in the input time series [23]. We used the software SPM1D, a parametric statistical testing method developed for nD time series [24], to control for FPR in the analysis of normalized joint moment profiles 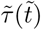, and quantify the effect of both factors (GS and SL) and of their interaction on the dependent variable in different phases of the gait cycle. 2-way repeated-measure ANOVA was conducted using the SPM1D software package, using SPM1D’s function anova2rm [24]. Inference was conducted setting a corrected type-I error rate *α* = 0.05 based on SPM1D’s correction based on random field theory (RFT) to estimate the smoothness in the input data.

After the main effect analysis, we conducted pairwise comparisons to establish the specific effect of SL on the measured joint moment profiles, testing for the null hypothesis that the mean profiles measured at the same speed for nominal and bio-feedback modulated SL conditions were not different from one another at any time point. This resulted in two comparisons (ss vs. increased SL, and ss vs. decreased SL) per speed, per joint, for a total of *n*_*comp*_ = 30 pairwise comparisons. Pairwise comparisons were conducted using two-tailed paired t-tests using SPM1D function `ttest_paired`, and defining thresholds t-scores for significance at *α* = 0.05 using a Bonferroni correction (*n* = *n*_*comp*_) on the paired difference thresholds calculated by SPM1D.

### Torque pulse approximation

We then conducted a second analysis with the specific purpose of composing a controller based on the application of pulses of torque at select phases of the gait cycle. As such, we sought to approximate the effect of SL modulation on joint moment profiles with a series of rectangular pulses of torque. First, the normalized difference 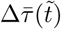 of subject specific average joint moment profiles 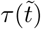 between conditions of positive or negative SL, and ss SL, at each gait speed *j* were extracted as:

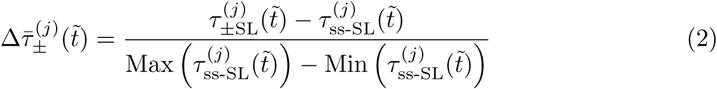

The rectangular torque pulses 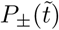 were defined using a constant duration of 10% of gait cycle, variable time of application *α*_*l*_, and amplitude *A*_*l*_ as in the equation:

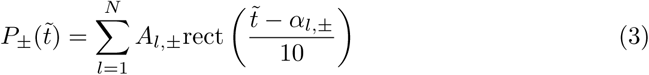

For this specific work, we examined the one and two pulse (*N* = 1, 2) approximation of the function 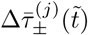, and used nonlinear constrained optimization—MATLAB function’s `fmincon`—to find the values of parameters *A* and *α* that minimize the norm of residuals 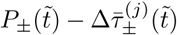. In this optimization, the domain for *α*_*l*_ was defined as the set of integers between 5% and 95%, providing a quantization in the time of application of torque pulses equal to 1% of a gait cycle duration. For each joint, we divided the estimated pulses into two different groups depending on whether their amplitude was in the positive, or negative direction (i.e., positive extension for the hip and knee joint, positive plantarflexion for the ankle joint).

With the purpose of identifying location and amplitude of application of pulses of torque that would approximate the SL-specific difference between joint moment profiles measured at all speeds, we performed statistical analyses to determine if any of the outcome measures, pulse magnitude and location, were significantly modulated by any of the three independent variables (i.e., joint, direction of SL modulation, and sign of applied pulse). For the purposes of our analysis, pulse magnitude is the absolute value of pulse amplitude. Four separate linear mixed effects models (SAS V9.4, SAS Institute, Cary, NC) were performed on the one and two pulse approximations for both the pulse magnitude and location data sets to test the null hypothesis that no independent variable had an effect on the outcome measures. The models included fixed effects for each of the independent variables as well as all two-way and one three-way interaction between them. Heterogeneity due to trials completed under different gait speed conditions, and multiple pulses in the case of the two pulse approximation were accounted for by the inclusion of random effects. Correlation between multiple measurements taken on the same subject were accounted for by the inclusion of a repeated measure effect. Upon comparing nested model Akaike information criterion (AIC) values, the lowest AIC value came from the unstructured covariance structure and was therefore selected for the final models.

In case of effects statistically significant at the *α* < 0.05 level, effects and interactions were further investigated through post hoc Tukey-Kramer tests (*α* = 0.05). The Tukey-Kramer post hoc test for the joint snd SL modulation interaction was used to establish the presence of significant differences in pulse magnitude means between joints for SL modulation conditions, separately. The three-way interaction between joint, SL modulation, and pulse sign was broken down using the Tukey-Kramer post hoc test to find the significant differences between torque pulse location, for a given joint, under different combinations of factors SL modulation and torque pulse sign. We were especially interested in testing whether there were specific instants of time where the application of a positive torque pulse would modulate SL in a certain direction, and, simultaneously, where the application of a negative torque pulse would modulate SL in the opposite direction. As such, for each joint, we conducted two pairwise comparisons: one to compare the location variables measured for positive pulse torque sign and positive SL modulation with the variables measured for negative pulse torque sign and negative SL modulation, and a second one to compare the location variables measured for positive pulse torque sign and negative SL modulation with the variables measured for negative pulse torque sign and positive SL modulation.

To visually illustrate the distributions of torque pulses in gait cycle, location histograms were generated. All pulses were combined across the twenty subjects and five gait speed conditions and grouped by joint and SL modulation condition for the one and two pulse approximations, separately. For representation purposes, within each histogram, the pulses are divided into positive and negative groups according to the sign of pulse amplitude 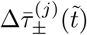, and then further divided into two more categories (i.e. small and large) based on whether their magnitude was smaller or larger than the group median.

## Results

### Protocol validation

The mean Froude numbers with 95% confidence intervals, across gait speed conditions, are shown in the left of Fig. 2. The use of the Froude did not reduce the across-subject variability in ss-GS, with *CV*_*Fr*_ = 0.106, slightly greater than *CV*_ss-GS_ = 0.104, and both smaller than 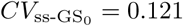. All differences account for an effect size that can be considered very small. The assessment of SL values across all conditions is shown in the center of Fig. 2. The change in mean SL from ss-SL across all ten feedback conditions for all subjects was equal to ±14.94%, close to the target ±17% value. The maximum standard deviation of SL_0_ values for all five non-feedback conditions, averaged across all subjects, was relatively small (*σ* _max_ = 3.86%). Based on these measures, we conclude that the protocol significantly modulated values of SL and GS, such that statistical analysis may be performed.

**Fig 2.**
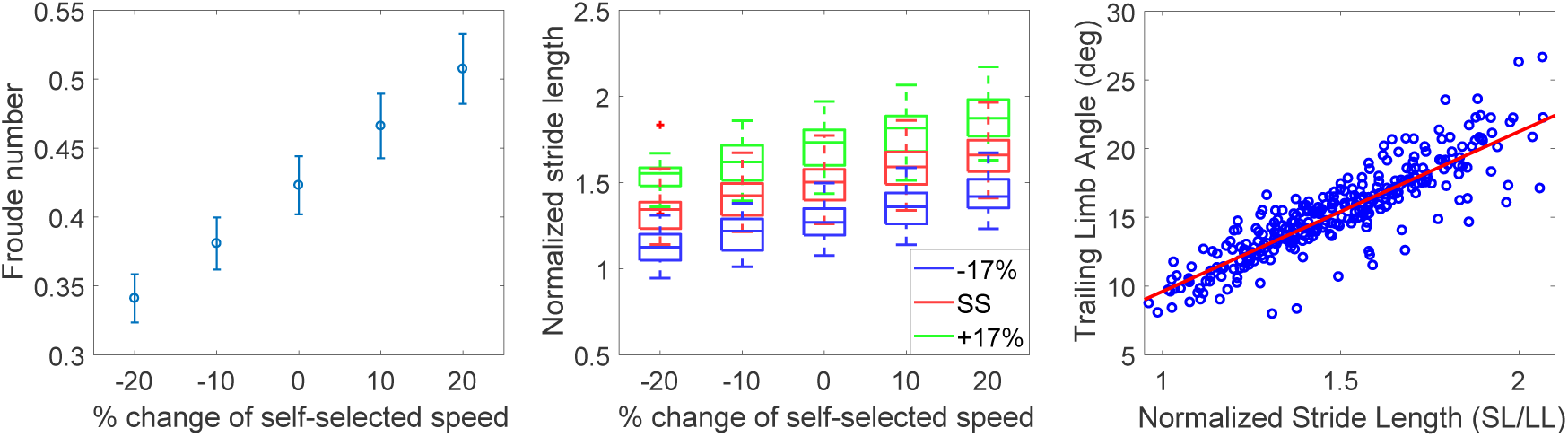
(Left) the distribution of mean Froude numbers with 95% confidence intervals at the various treadmill imposed GSs, (center) normalized stride lengths measured at various speeds and biofeedback conditions. The box plot shows the median as a horizontal line, and the box at 25% and 75% percentiles, with whiskers extending to ± 3*σ,* and (right) mean trailing limb angle and mean normalized stride length for each of the 15 conditions for each of the 20 subjects. Linear regression indicates that there is a strong correlation (r = 0.87) between the two measures.

As seen in the right side of Fig. 2, linear regression demonstrated a strong correlation (r = 0.87) between SL_0_ and TLA, which indicates that subjects also modulated TLA while achieving biofeedback cued SL modulation. The group analysis of joint moments in the three SL conditions across five GS conditions is depicted in Fig. 3

**Fig 3.**
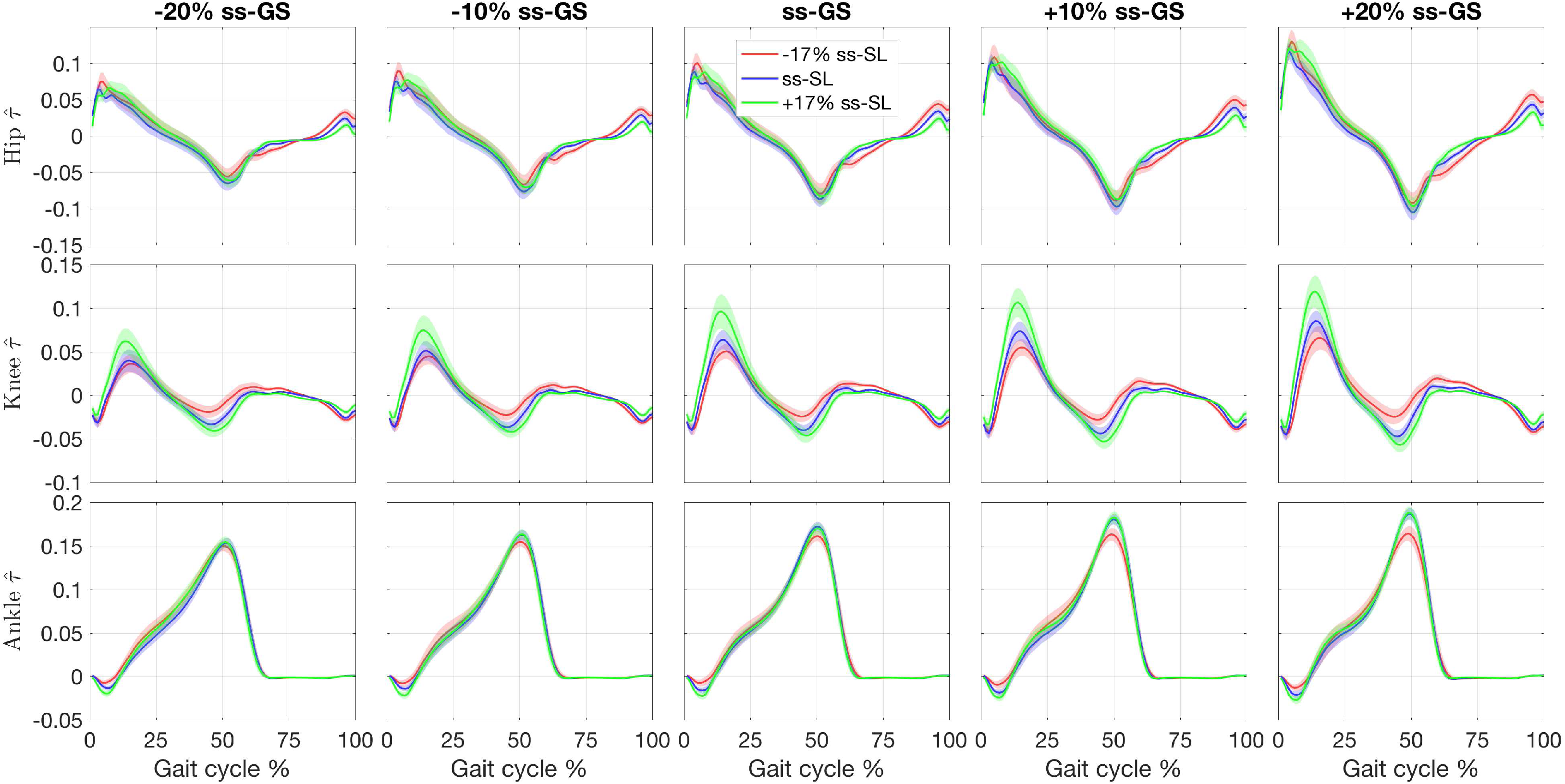
Effect of gait speed (GS) and stride length (SL) modulation on the normalized joint moments 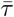. Joints are organized by row, GS are organized by columns, relative to the subject-specific ss-GS. Conditions corresponding to cued SL values are superimposed on each plot. Lines indicate the group mean, with the shaded region indicating the standard error.

### Continuum analysis

The continuum analysis showed an effect of GS on the normalized joint moment profiles, where a significant effect of GS was detected for a total 82.4%, 80.4% and 64.7% of the gait cycle for the hip, knee, and ankle joint respectively, as shown in Fig. 4. The effect of SL on hip joint moment was significant for seven short clusters in early to midstance, at push-off, and two clusters spanning the majority of swing for a total duration of a significant effect of SL on hip joint moment of 56.1% of the gait cycle. A stronger effect, both in magnitude and duration, was measured at the knee joint with the first two clusters spanning early stance, and continuing in the swing phase. The effect of SL on knee joint moment was highly significant from midstance until the end of gait cycle; for a total of 91.2% of the gait cycle with a significant effect. A strong effect of SL was detected at the ankle joint for 5 clusters; at weight acceptance, push-off and three clusters covering approximately half of swing for a total of 41.7% of the gait cycle. The interaction between the two factors was significant for the hip for four clusters; mainly during the transition from push-off and during late swing for a total of 33.9% of the gait cycle. For the knee, the interaction was significant for four clusters; during early stance, late stance, early swing, and midswing for a total of 35.9% of gait cycle. For the ankle, the interaction was only significant for two clusters, late stance and mid swing for a total of 19.1% of gait cycle. Pairwise comparisons of joint moment profiles measured at nominal and biofeedback-modulated SL values are shown in Fig. 5 – 7 for all GSs.

**Fig 4.**
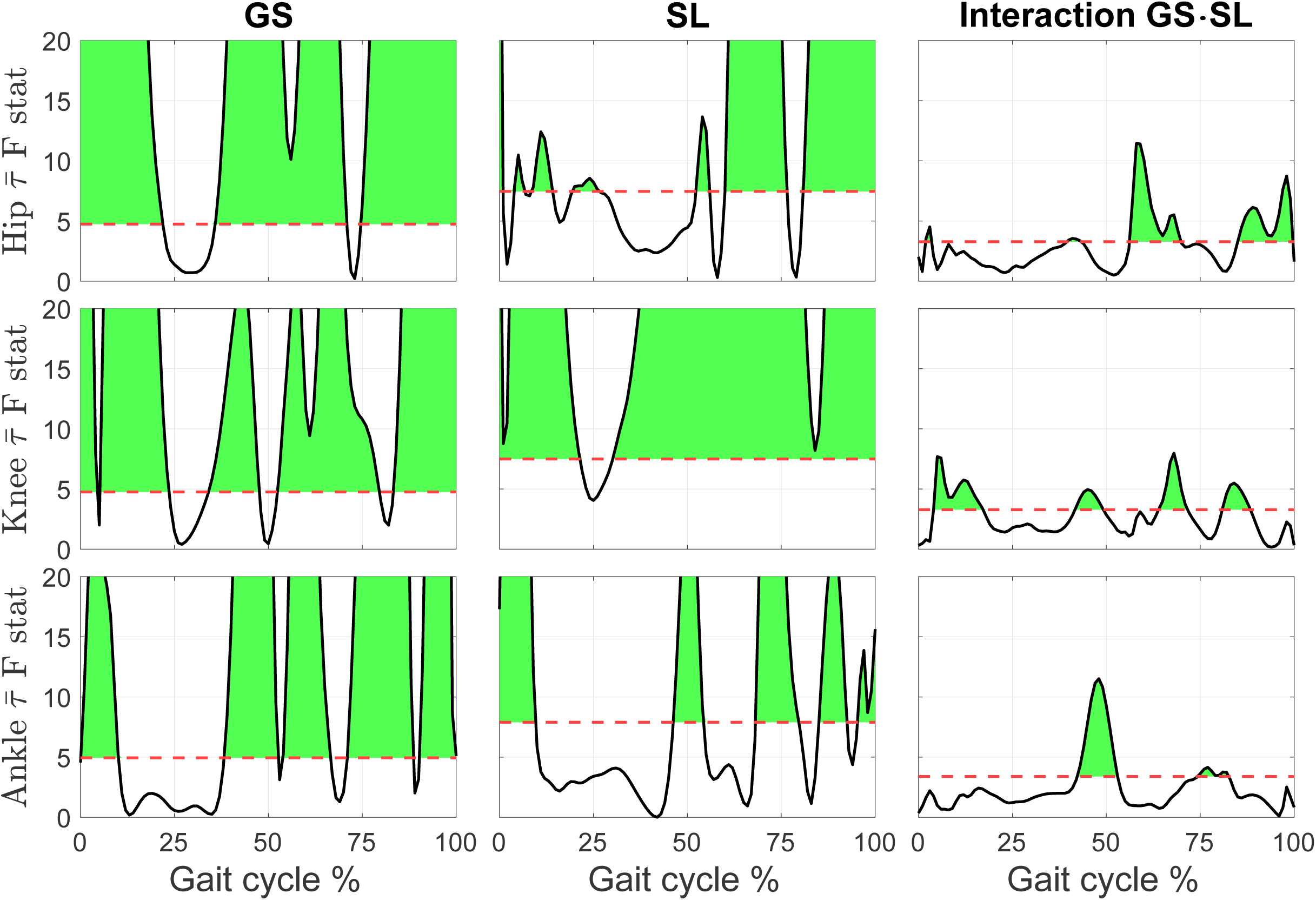
Main effects of gait speed (GS) and stride length (SL), and of their interaction, on the normalized joint moment profiles during normal walking, as described by the 1D time series of F-statistic extracted by the 2-way repeated measure ANOVA. The threshold F statistic for each experimental condition is reported by the red dashed line, and values above (shaded in green) correspond to a significant group effect of the factor, at the corresponding gait cycle instant, for a corrected type I error rate *α* < 0.05.

**Fig 5.**
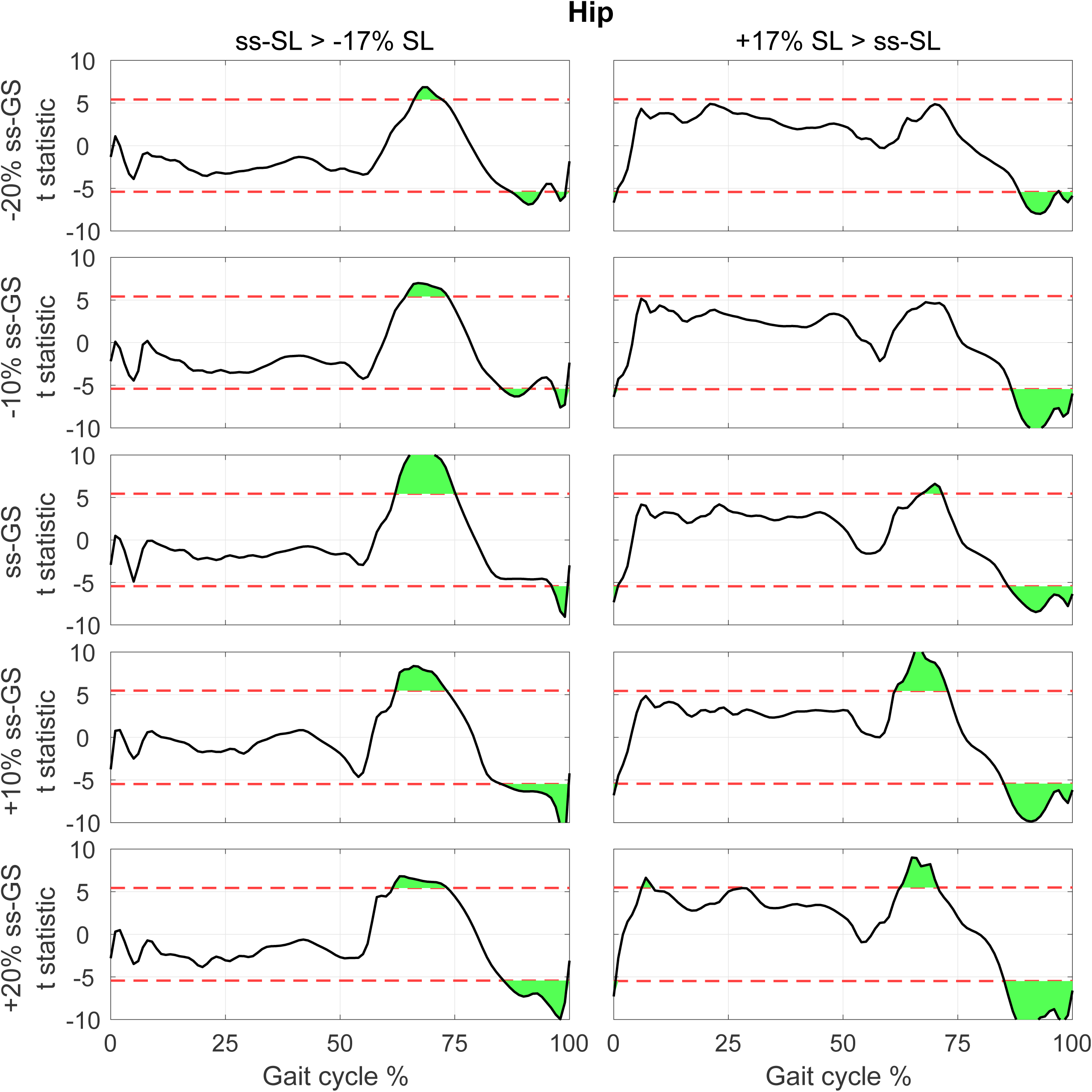
T-statistic resulting from pairwise comparisons of normalized hip torque moment at normal and modulated SL (columns), measured at the same GS, for each GS (row). Red dashed lines show the threshold *t* value that provides a corrected type I error rate *α* = 0.05, extracted using a Bonferroni correction that accounts for all pairwise comparisons *n*_*comp*_ = 30.

Pairwise comparisons for the hip joint show that during increased SL conditions (Fig. 5, right), hip flexion moment during early swing and hip extension moment during late swing decreased. The first comparison reached significance in three out of five GS conditions, while the second effect was significant at all GS values. No effect of SL on hip joint moment during stance were observed in more than one GS condition. A similar pattern was observed when SL is decreased via biofeedback (Fig. 5, left).

For the knee, during increased SL conditions (Fig. 6, right), knee extension moment increased in early stance, while knee flexion moment increased in late stance. During the swing phase, knee extension moment decreased in early swing, and knee flexion moment decreased in late swing. The effects reported were significant at the group level at all GSs. A similar pattern was observed for a decrease of SL, with smaller effects for the increased knee extension at early stance (a significant effect was measured only in four out of five GS conditions, Fig. 6, right).

**Fig 6.**
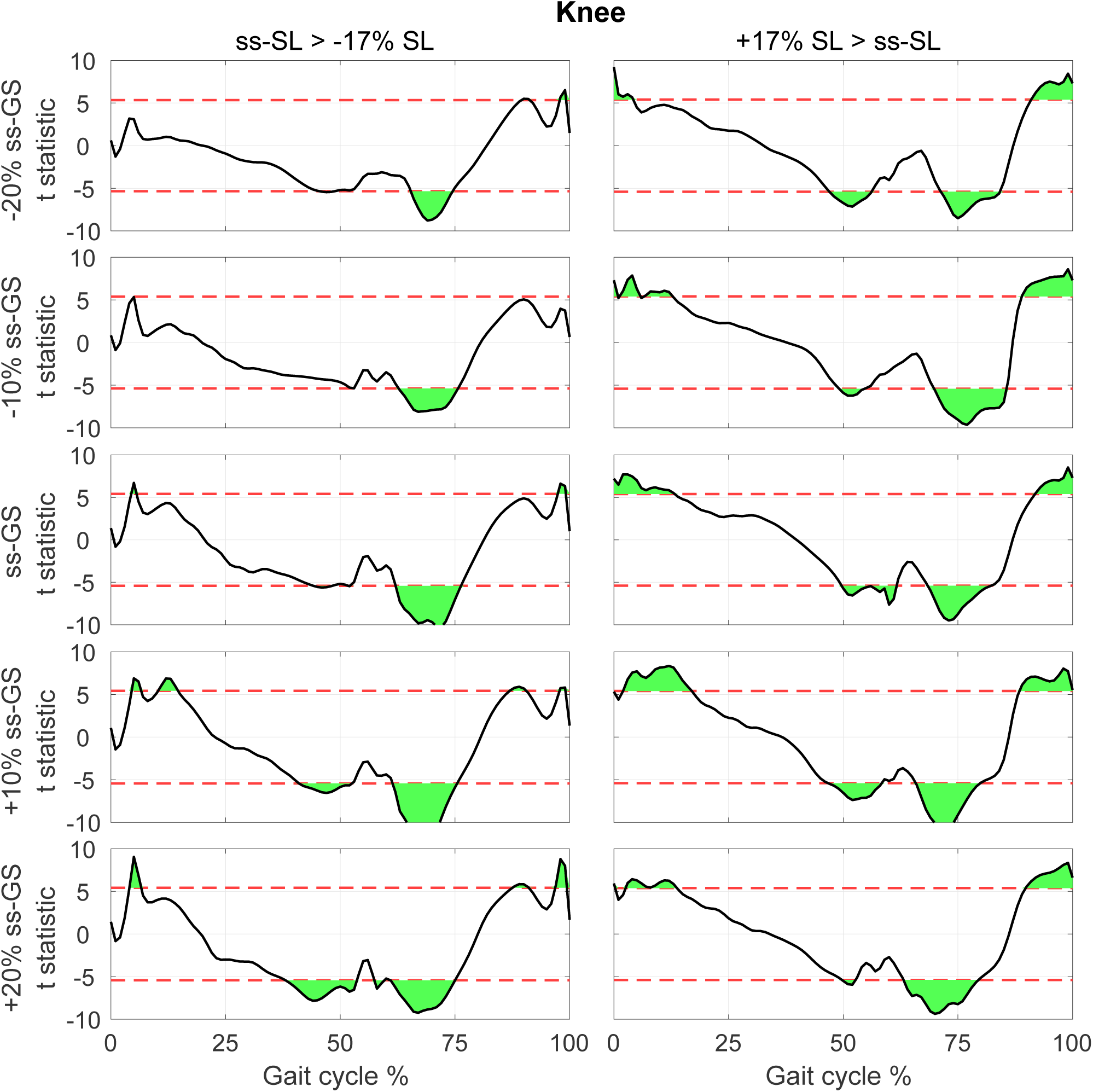
T-statistic resulting from pairwise comparisons of normalized knee torque moment at normal and modulated SL (columns), measured at the same GS, for each GS (row). Red dashed lines show the threshold *t* value that provides a corrected type I error rate *α* = 0.05, extracted using a Bonferroni correction that accounts for all pairwise comparisons *n*_*comp*_ = 30.

For the ankle, during increased SL conditions (Fig. 7, right), ankle dorsiflexion moment increased at early stance, while no effect on plantarflexion moment was measured at push-off. A similar pattern was observed for a decrease of SL (Fig. 7, right), with a greater effect measured in terms of increased plantarflexion moment at early swing (significant at all GSs). In two out of five GS conditions, the decreased SL condition exhibited a reduced ankle plantarflexion moment during push-off significant at the group level.

**Fig 7.**
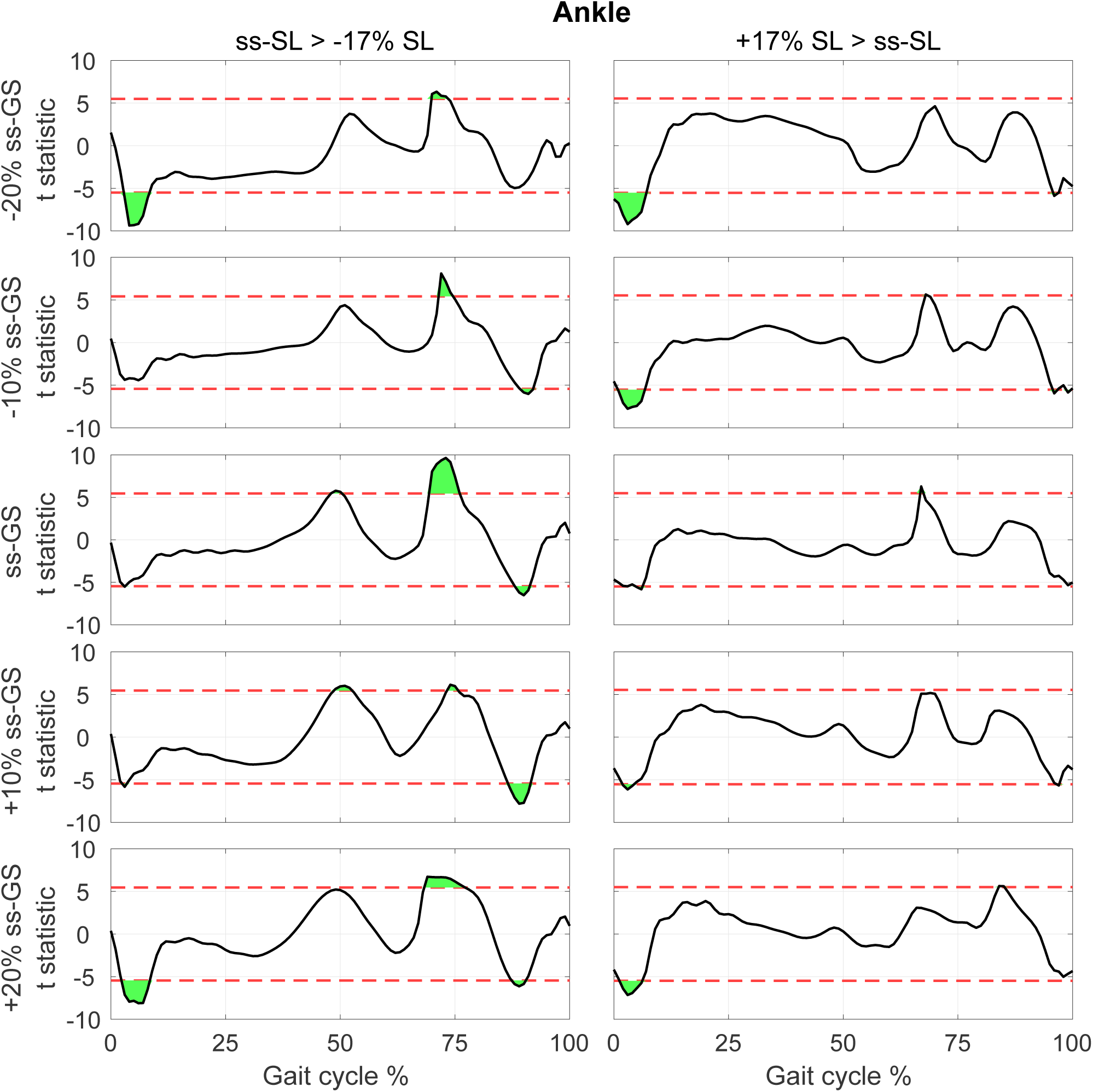
T-statistic resulting from pairwise comparisons of normalized ankle torque moment at normal and modulated SL (columns), measured at the same GS, for each GS (row). Red dashed lines show the threshold *t* value that provides a corrected type I error rate *α* = 0.05, extracted using a Bonferroni correction that accounts for all pairwise comparisons *n*_*comp*_ = 30.

### Torque pulse approximation

Figures 8 and 9 show the distribution of torque pulse magnitudes grouped by joint for both positive and negative SL modulations for the one and two pulse approximations, respectively.

**Fig 8.**
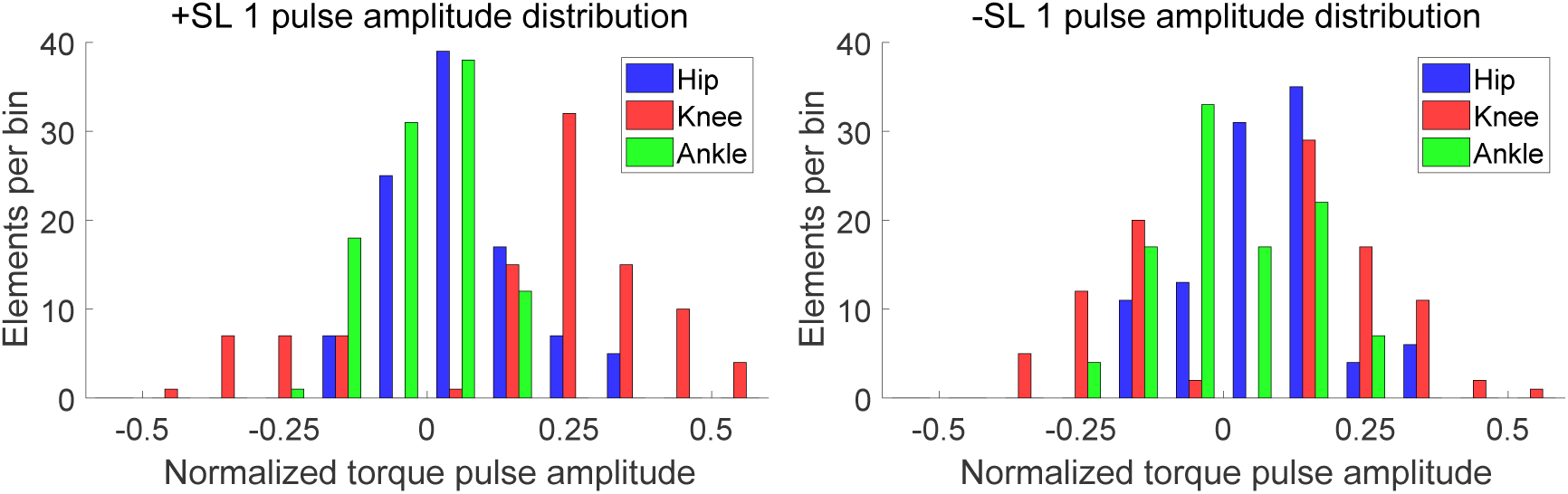
Histogram of one pulse approximation normalized amplitudes, sorted by SL modulation and joint.

**Fig 9.**
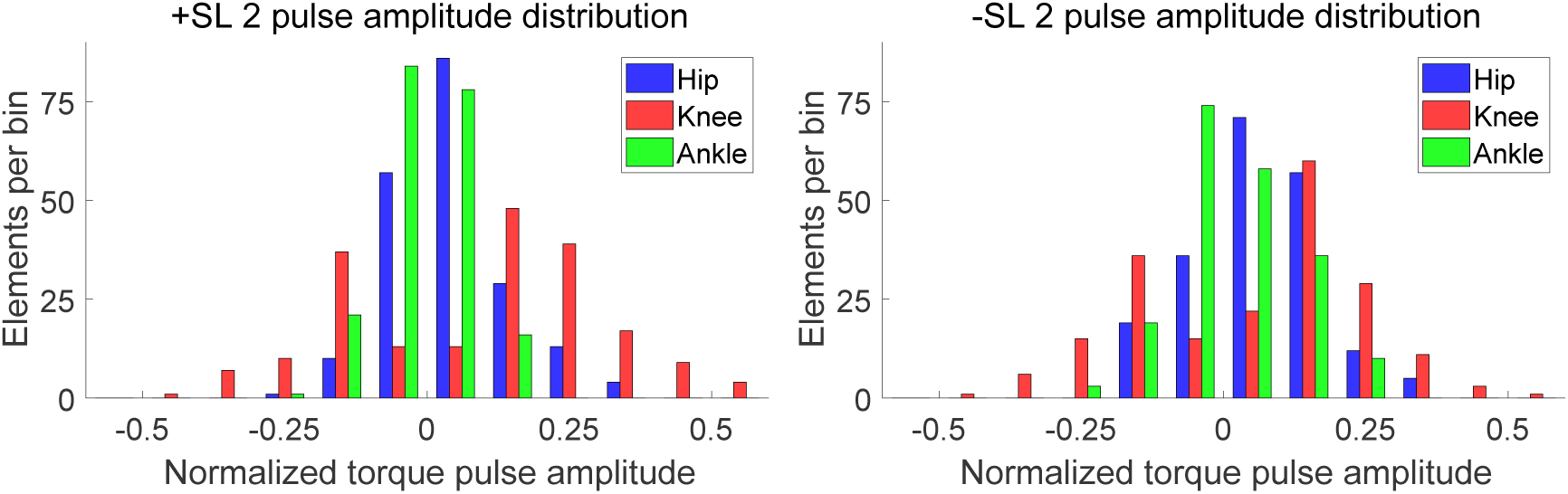
Histogram of two pulse approximation normalized amplitudes, sorted by SL modulation and joint.

The results of the linear mixed effects model analyses for torque pulse magnitude are shown in Tables 1 and 2. A significant effect of factors joint and pulse sign, and a significant interaction between factors joint and SL modulation were observed for the one and two pulse approximations. The interaction between factors joint, SL modulation, and pulse sign was significant for the two pulse approximation. The Tukey-Kramer post hoc test for the joint and SL modulation interaction is shown in Tables 3 and 4. The mean and standard deviation of the joint magnitudes are shown in Figures 10 and 11. In the one and two pulse approximations, for both SL modulation conditions, the normalized torque pulse magnitudes of the knee joint were greater than both the hip and ankle joints. The only significant difference between hip and ankle joint magnitudes existed for the positive SL modulation condition for the two pulse approximation.

**Table 1.** Magnitude linear mixed effects model results for the one torque pulse approximation

| Factor | DF | F Value | Prob >F |
| --- | --- | --- | --- |
| Joint | 2 | 130.63 | <0.001 |
| SL Mod | 1 | 0.50 | 0.482 |
| Pulse Sign | 1 | 5.44 | 0.020 |
| Joint x SL Mod | 2 | 12.41 | <0.001 |
| Joint x Pulse Sign | 2 | 1.15 | 0.318 |
| SL Mod x Pulse Sign | 1 | 0.03 | 0.862 |
| Joint x SL Mod x Pulse Sign | 2 | 2.14 | 0.120 |

**Table 2.** Magnitude linear mixed effects model results for the two torque pulse approximation

| Factor | DF | F Value | Prob >F |
| --- | --- | --- | --- |
| Joint | 2 | 154.81 | <0.001 |
| SL Mod | 1 | 0.21 | 0.645 |
| Pulse Sign | 1 | 24.68 | <0.001 |
| Joint x SL Mod | 2 | 10.75 | <0.001 |
| Joint x Pulse Sign | 2 | 2.91 | 0.055 |
| SL Mod x Pulse Sign | 1 | 2.05 | 0.152 |
| Joint x SL Mod x Pulse Sign | 2 | 3.99 | 0.019 |

**Table 3.** Magnitude Tukey-Kramer post hoc test results for the one torque pulse approximation

|  | Mean (Std Err Dif) |  |  |
| --- | --- | --- | --- |
| SL Mod | Hip - Knee | Hip - Ankle | Knee - Ankle |
| +SL | -0.172 (0.014) | 0.022 (0.008) | 0.194 (0.014) |
| -SL | -0.094 (0.011) | 0.010 (0.010) | 0.104 (0.012) |

**Table 4.** Magnitude Tukey-Kramer post hoc test results for the two torque pulse approximation

|  | Mean (Std Err Dif) |  |  |
| --- | --- | --- | --- |
| SL Mod | Hip - Knee | Hip - Ankle | Knee - Ankle |
| +SL | -0.115 (0.009) | 0.019 (0.005) | 0.135 (0.009) |
| -SL | -0.069 (0.007) | 0.015 (0.006) | 0.085 (0.008) |

**Fig 10.**
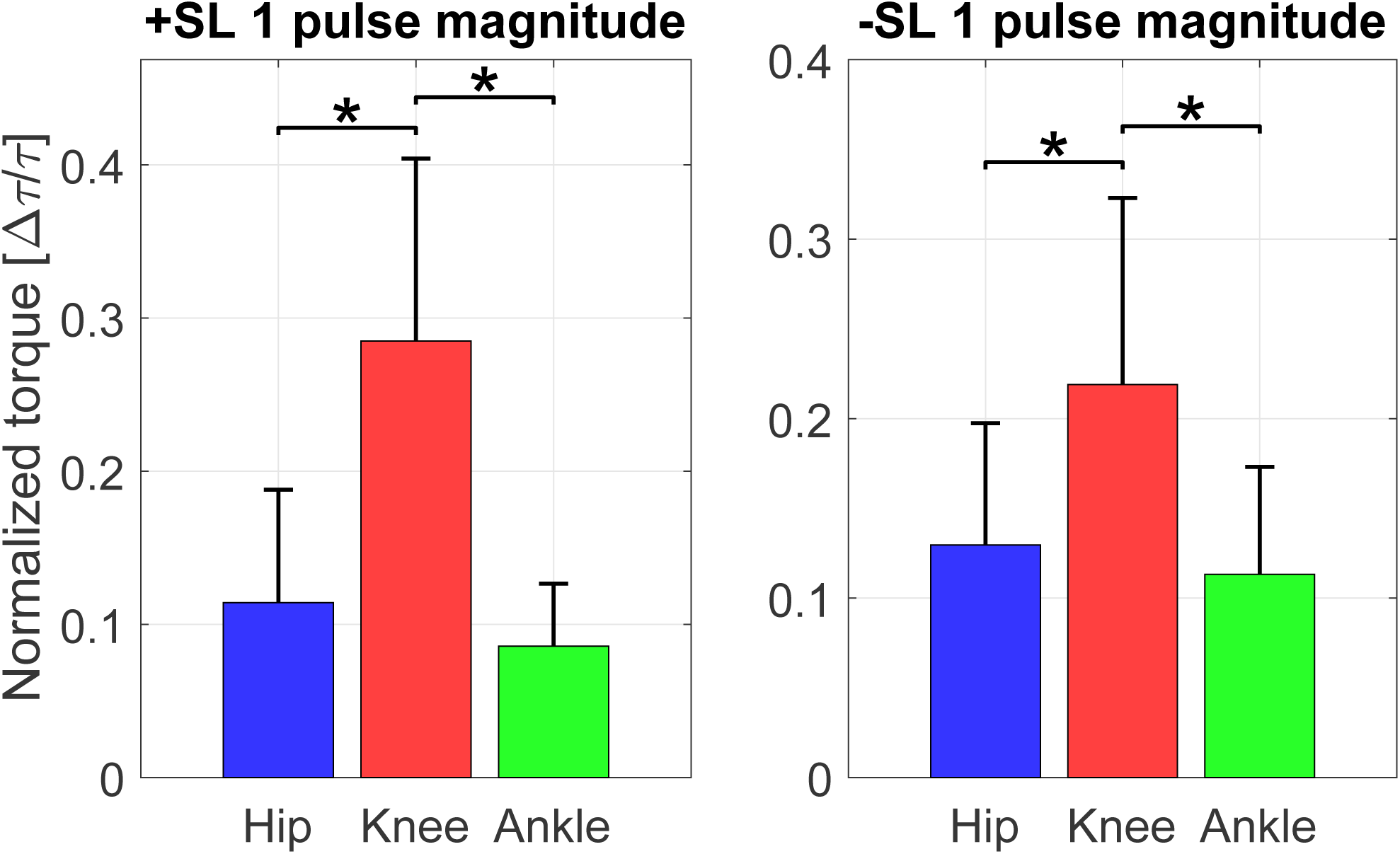
Pulse magnitude by joint and SL modulation for the one pulse approximation (mean ± standard deviation). Asterisks indicate pairwise comparisons significant at the *p* < 0.05 corrected level.

**Fig 11.**
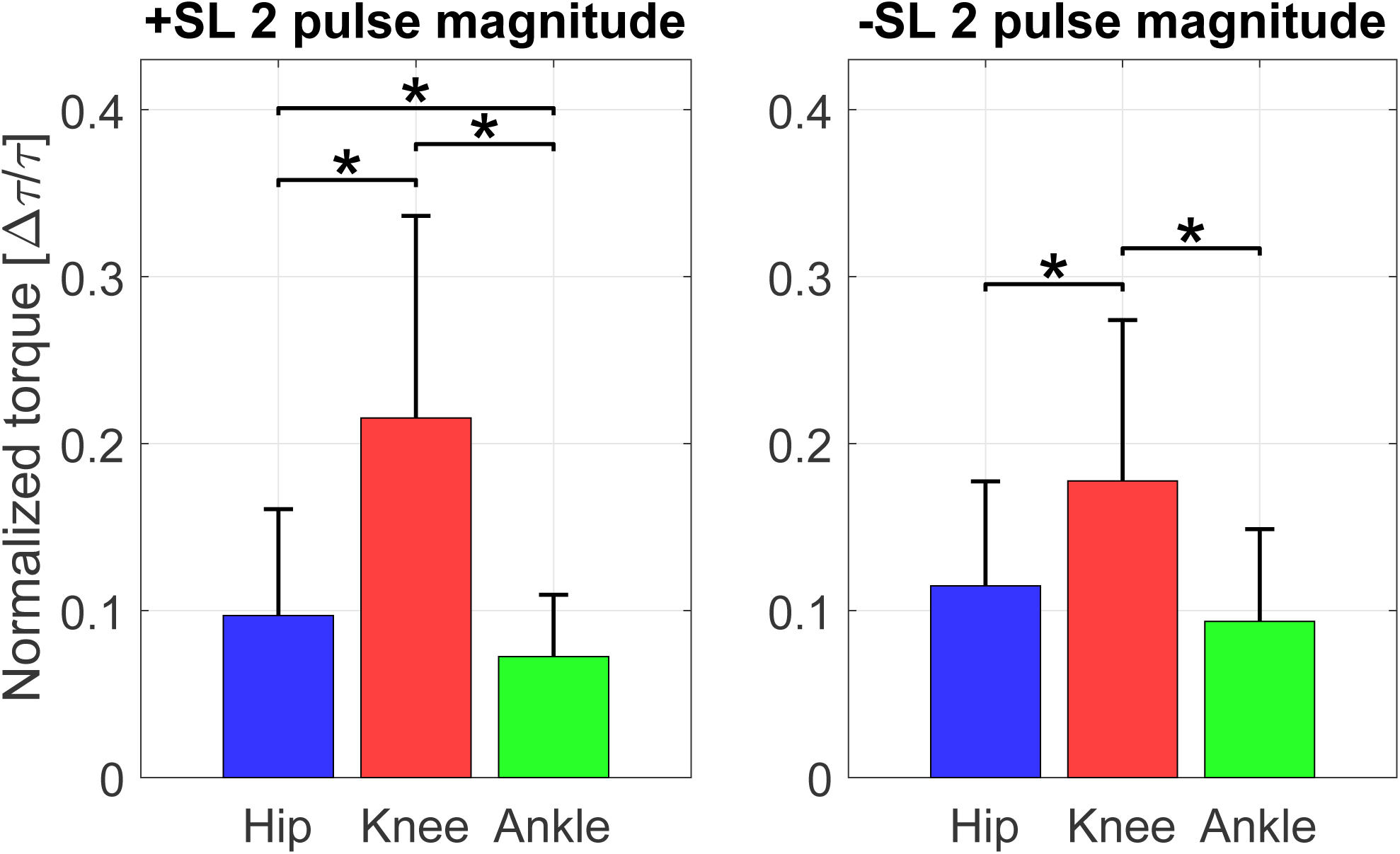
Pulse magnitude by joint and SL modulation for the two pulse approximation (mean ± standard deviation). Asterisks indicate pairwise comparisons significant at the *p* < 0.05 corrected level.

Figures 12 and 13 show the distributions of torque pulse location in gait cycle.

**Fig 12.**
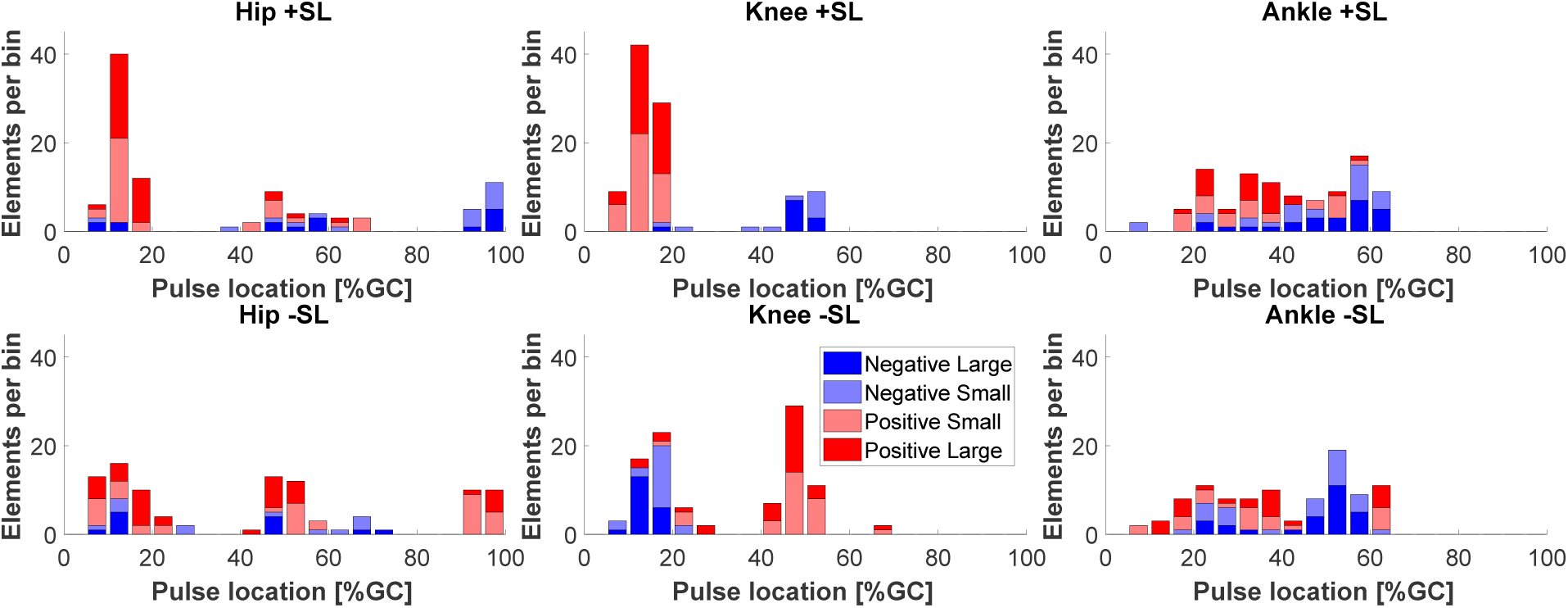
Histogram of one pulse approximation locations in gait cycle, sorted by pulse amplitude sign and magnitude.

**Fig 13.**
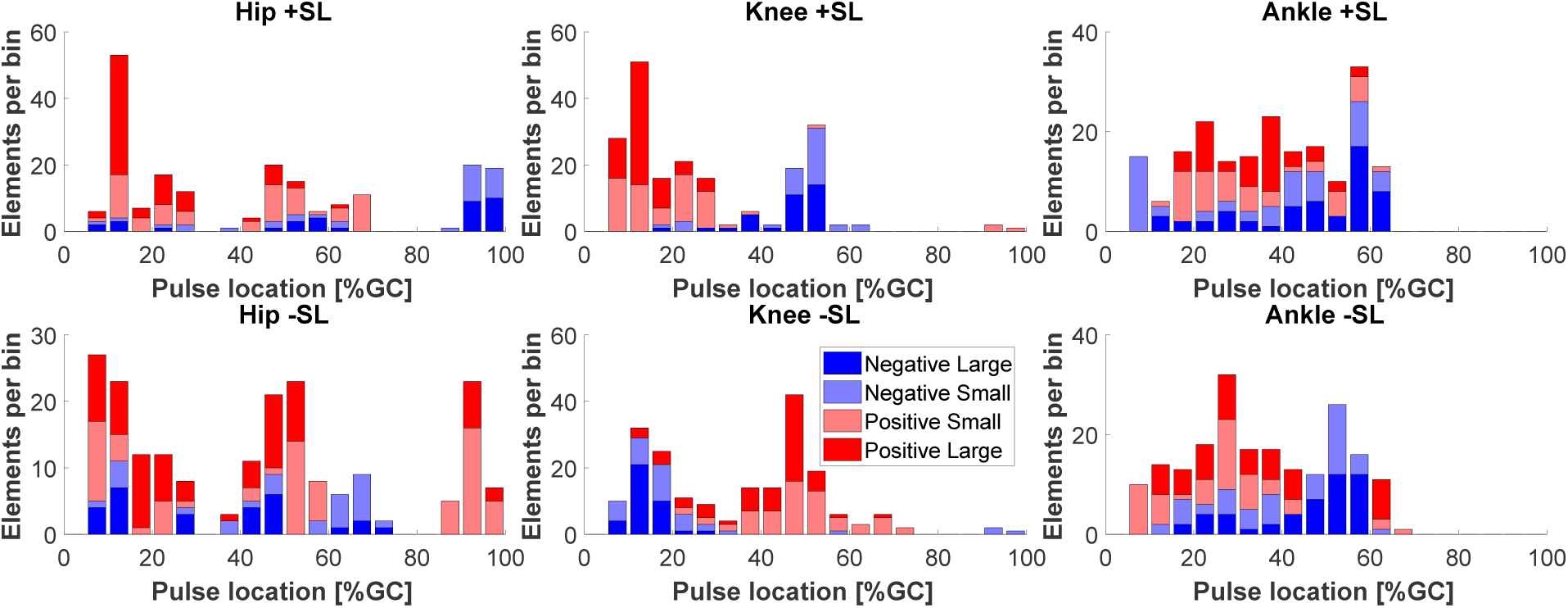
Histogram of two pulse approximation locations in gait cycle, sorted by pulse amplitude sign and magnitude.

The linear mixed effects model results for torque pulse location in gait cycle are shown in Tables 5 and 6. The analyses yielded a highly significant effect of joint and interactions between joint and pulse sign, SL modulation and pulse sign, and joint, SL modulation, and pulse sign for both pulse approximations. The factor of pulse sign was only significant for the one pulse approximation and the factor of SL modulation and interaction of joint and SL modulation were only significant for the two pulse approximation.

**Table 5.** Location linear mixed effects model results for the one torque pulse approximation

| **Factor** | **DF** | **F Value** | **Prob >F** |
| --- | --- | --- | --- |
| Joint | 2 | 49.32 | **<0.001** |
| SL Mod | 1 | 2.14 | 0.146 |
| Pulse Sign | 1 | 31.48 | **<0.001** |
| Joint x SL Mod | 2 | 0.53 | 0.589 |
| Joint x Pulse Sign | 2 | 13.61 | **<0.001** |
| SL Mod x Pulse Sign | 1 | 166.41 | **<0.001** |
| Joint x SL Mod x Pulse Sign | 2 | 78.42 | **<0.001** |

**Table 6.**
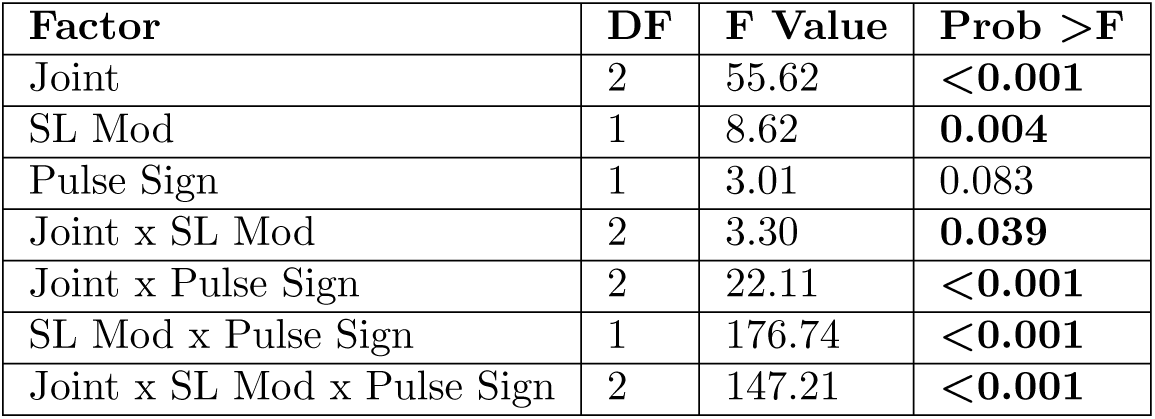
Location linear mixed effects model results for the two torque pulse approximation

All of the pairwise comparisons conducted to assess the symmetry of the effect for a reversal of SL conditions are reported in Tables 7 and 8. In the one pulse approximation, both pairwise comparisons for the hip and ankle joint yielded a relatively large differences (9 - 11% gait cycle duration) in mean location, with three out of the four comparisons statistically significant. On the contrary, the pairwise comparisons for the knee yielded small (1 - 2% gait cycle duration) and statistically insignificant mean differences. This indicates that for the knee joint, clustering of torque pulses by location was symmetrical in reversed stride length conditions, with negative pulses in negative SL conditions clustering around a similar value as positive pulses in positive SL conditions, and positive pulses in negative SL conditions clustering around a similar value as negative pulses in positive SL conditions, while the same effect was not measured for the hip and ankle joints. However, this pattern was not observed in the two pulse approximation; in which one out of the four hip and ankle joint comparisons and one of the two knee joint comparisons were statistically significant. For the one and two pulse approximations, all knee joint mean comparisons were below 10% gait cycle, the width of the torque pulses used for the approximation.

**Table 7.** Location Tukey-Kramer post hoc testing for the one torque pulse approximation

|  | Mean[1] - Mean[2] (Std Err Dif) |  |
| --- | --- | --- |
| Joint | [1] +SL +Sign<br>[2] -SL - Sign | [1] - SL + Sign<br>[2] +SL -Sign |
| Hip | -9 (7) | **-19 (5)** |
| Knee | -1 (2) | -2 (2) |
| Ankle | **-11 (3)** | **-13 (3)** |

**Table 8.** Location Tukey-Kramer post hoc testing for the two torque pulse approximation

|  | Mean[1] - Mean[2] (Std Err Dif) |  |
| --- | --- | --- |
| Joint | [1] +SL +Sign<br>[2] -SL - Sign | [1] - SL + Sign<br>[2] +SL -Sign |
| Hip | -3 (4) | **-20 (4)** |
| Knee | **8 (2)** | 6 (2) |
| Ankle | -1 (2) | -3 (2) |

## Discussion

We exposed subjects to a factorial modulation of gait speed (GS) and stride length (SL) and utilized inverse dynamics to estimate the lower extremity joint moments in the sagittal plane. With our protocol, we modulated SL of individuals significantly between conditions, with a mean change in SL equal to ±15% of the self-selected value, close to the target ±17%. Furthermore, inter-individual variability in self-selected gait speed (ss-GS) was reasonably small, with a coefficient of variation *CV*_ss-GS_ = 0.103. Based on these measures, it is apparent that our protocol significantly modulated both SL and GS, such that statistical analysis may be performed to assess changes in joint kinetics arising from exposure to these conditions. Our analysis showed a strong correlation (r = 0.87) between SL and TLA, indicating that TLA was indirectly modulated through the explicit cueing of SL modulation. Our data analyses focused primarily on the effects on joint kinetics introduced by modulation of SL at various GSs and secondarily on the effects introduced by GS.

### Knee joint moment

The most prominent effects of SL modulation on joint moment were observed for the knee joint. Simple visual inspection of the normalized joint moment profiles, Fig. 3, clearly indicates the effect of SL on joint moment, where stance phase peak extension moment and peak flexion moment increase with increasing SL as well as increasing GS. Increasing peak knee extension moment with SL was also observed previously [13, 18] while increasing peak flexion moment is in agreement with one previous study [13] but also in contrast with previously observed decreasing peak knee flexion moment [18]. This contrasting result could be attributed to important differences with the experimental paradigm pursued in [18], where SL and cadence, and not SL and gait speed, were cued. These observations are in agreement with the continuum analysis, where a main effect of SL was observed for 91.2% of the gait cycle, the highest percentage of all three joints. The significant effect of SL on peak flexion and extension moments during stance is supported by the pairwise comparisons between joint torque measured at different SL conditions (Fig. 6). Here, significant effects at early stance support an increase in knee extension moment with increasing SL, and significant effects in late stance support an increase in knee flexion moment with increasing SL. These effects introduced by increases in SL are also captured by the torque pulse approximations through visual inspection of the pulse approximation histograms (Fig. 12 and 13), and the findings of their associated linear mixed effects model pulse location analyses. Our findings indicate that an increase in SL is associated with positive pulses of torque – an increase in extension moment – in early stance, and negative pulses – an increase in flexion moment – in late stance. The reverse pattern is observed for SL decrease, where negative pulses of torque – decreasing extension moment – are extracted in early stance, and positive pulses of torque – decreasing flexion moment – in late stance. This pattern is supported by the Tukey-Kramer post hoc one pulse approximation results for the joint, SL modulation, and pulse sign interaction effect of the pulse location linear mixed effects model analyses. Furthermore, for the one pulse approximation, there is no significant difference in location between pulses of positive SL and negative sign and pulses of negative SL and positive sign. The lack of significant difference in the location of these specific pulse groups supports the observation of a systematic pulse pattern at the knee which reverses in sign with reversal of SL modulation direction. Another indication of the effect of SL modulation on knee joint moment derives from the fact that the one and two pulse approximations yielded a significant effect of the joint and SL modulation interaction on magnitude (*p <* 0.001) with Tukey-Kramer post hoc tests indicating the knee having the greatest normalized pulse magnitude (Tables 3 and 4 and Figures 10 and 11). Overall, our findings indicate that the effects of SL on the knee joint moment can be described as follows: knee extension moment at early stance and knee flexion moment at late stance are increased for an increase in SL, while knee extension moment at early stance and knee flexion moment at late stance are decreased for a decrease in SL.

### Hip joint moment

Significant effects of SL modulation were also observed at the hip joint. As indicated by the main effect of SL quantified by the continuum analysis, there are two major intervals of significance: early swing and late swing. These effects in swing are visible upon inspection of the group moment profiles where an increase in SL is associated with a decrease in flexion moment during early swing and a decrease in extension moment during late swing. These findings are in contrast with a previous study which only observed a decrease in peak hip flexion moment [18]. The statistical significance of these observations is clearly indicated by the pairwise comparisons of SL modulation at different speeds shown in Fig. 5. Here, in all pairwise comparisons, an increase in SL is associated with a significant interval in early swing and in late swing. Another indication on how SL modulates hip moment during the swing phase is provided by the torque pulse approximation histograms, which consistently depict a grouping of negative pulses in late swing for positive change in SL and positive pulses in late swing for negative change in SL. However, the pattern is less clear than the one seen at the knee joint because of the small magnitude of those pulses occurring in the swing phase, which even though they are representative of a statistically significant effect, they account for a small amplitude (see distribution of larger pulses in Fig. 12 and 13). As such, the effect in SL obtained for a change in magnitude of the applied torque is not symmetrical (Tables 7 and 8), as demonstrated by the Tukey-Kramer post hoc tests for both pulse approximations. These groupings of pulses in late swing are likely associated with the decrease in hip extension moment during late swing associated with increasing SL.

### Ankle joint moment

Relatively small effects of SL modulation were observed at the ankle joint. Most prominently, during loading response, dorsiflexion moment increases with an increase in SL as also can be observed in the group moment profiles. This is supported by the main effect of SL measured via the continuum analysis, where the first 10% of gait cycle shows a significant effect of GS on joint moment. This is further supported by the observation of a significant increase in dorsiflexion moment (negative change) for increases in SL, in nine of the ten pairwise comparisons at all speeds, shown in Fig. 7. Another observed effect through the pairwise comparisons conducted via the continuum analysis is the increase in plantarflexion moment at early swing with increasing SL. This effect can be confirmed through visual inspection of the group moment profiles. In contrast with previous work [18], we did not observe a consistent increase in peak plantarflexion moment with increasing SL, with only the three highest speed conditions showing a significant effect for the transition from −17% ss to ss-SL, and no significant effects measured for an increase in stride length over the self-selected value. Also, increasing ankle plantarflexion moment with stride length at 40% of stance was not observed, in contrast with previous work [18].

## Conclusion

Our study has measured the effects of stride length (SL) on the lower extremity joint moment profiles at different speeds, demonstrating several consistent effects in our population. The main effects of increasing SL at the knee include an increase in knee extension moment at early stance and an increase in flexion moment at late stance. At the hip, the main effects of increasing SL are a decrease in flexion moment during early swing and a decrease in extension moment during late swing. For an increase in SL, the ankle primarily exhibits an increase in dorsiflexion moment during loading response. Given the observed linear relationship between SL_0_ and TLA, pulse torque approximation patterns associated with SL modulation are also associated with TLA modulation. These findings suggest that a possible joint moment assistance strategy based on pulses of torque applied primarily at the hip and the knee joint could induce modulations in both SL and TLA. According to our analysis, the application of an extension pulse of torque in early stance and a flexion pulse in late stance to the knee appear to be suitable assistance strategies to support an increase of SL and TLA during walking. If pulse torque assistance is to be applied at an additional joint, pulse torque assistance could be applied at the hip with a flexion pulse applied during late swing.

This study has some limitations. The methods pursued in this paper are based on a group analysis of joint moment profiles measured via inverse-dynamics. As such, it is possible that the most successful assistance strategies may significantly change between different individuals. Therefore, the group analysis based assistance strategy candidate could be best utilized as an initial estimate and assistance strategies could be iteratively optimized for each subject using human-in-the-loop optimization, like it has been done for single-joint assistance schemes [10].

Moreover, the proposed assistance strategy is based on the assumption that human contribution will not change when an assistive torque is applied via a wearable exoskeleton, such that the combination of torques applied by the two agents would result in a simple summation. However, it is well known that the human neuromuscular system is non-linear [25] and involves multiple complex feedback loops [26]. As such, the response to a torque perturbation at a specific instant in gait cycle will be difficult to predict. Given the difficulty of formulating a model of the human response to these assistance strategies, once again the results of this analysis work could be used as an initial estimate to be iteratively optimized for each subject using human-in-the-loop optimization.

## Acknowledgments

Research project funded by NSF-NRI-1638007, Instrumentation provided by NIH-GM-103333, NSF-REU-1460757, and NIH-P20GM109090.

## References

1. Mehrholz J, Thomas S, Werner C, Kugler J, Pohl M, Elsner B. Electromechanical-assisted training for walking after stroke (Review). Cochrane Database of Systematic Reviews. 2017;(5). doi:10.1002/14651858.CD006185.pub4. www.cochranelibrary.com.

2. Holanda LJ, Silva PMM, Amorim TC, Lacerda MO, Simão CR, Morya E. Robotic assisted gait as a tool for rehabilitation of individuals with spinal cord injury: A systematic review. Journal of NeuroEngineering and Rehabilitation. 2017;14(1):1–7. doi:10.1186/s12984-017-0338-7.

3. Schmid A, Duncan PW, Studenski S, Lai SM, Richards L, Perera S, et al. Improvements in Speed-Based Gait Classifications Are Meaningful. Stroke. 2007;38:2096–2101. doi:10.1161/STROKEAHA.106.475921.

4. Middleton A, Fritz SL, Lusardi M. Walking speed: The functional vital sign. J Aging Phys Act. 2015;23(2):314–322. doi:10.1123/japa.2013-0236.Walking.

5. Peterson CL, Cheng J, Kautz SA, Neptune RR. Leg extension is an important predictor of paretic leg propulsion in hemiparetic walking. Gait & Posture. 2010;32(4):451–456. doi:10.1016/j.gaitpost.2010.06.014.

6. Hsiao H, Knarr BA, Higginson JS, Binder-Macleod SA. Mechanisms to increase propulsive force for individuals poststroke. Journal of Neuroengineering and Rehabilitation. 2015;12(40). doi:10.1186/s12984-015-0030-8.

7. Hsiao H, Knarr BA, Higginson JS, Binder-Macleod SA. The Relative Contribution of Ankle Moment and Trailing Limb Angle to Propulsive Force during Gait. Human Movement Science. 2015; p. 212–221. doi:10.1016/j.humov.2014.11.008.

8. Browne MG, Franz JR. More push from your push-off: Joint-level modifications to modulate propulsive forces in old age. Plos One. 2018;13(8):e0201407. doi:10.1371/journal.pone.0201407.

9. Ahn J, Hogan N. Walking is Not Like Reaching: Evidence from Periodic Mechanical Perturbations. PLoS ONE. 2012;7(3). doi:10.1371/journal.pone.0031767.

10. Zhang J, Fiers P, Witte KA, Jackson RW, Poggensee KL, Atkeson CG, et al. Human-in-the-loop optimization of exoskeleton assistance walking. Science. 2017;356(June):1280–1284. doi:10.1126/science.aal5054.

11. Awad LN, Bae J, O’Donnell K, De Rossi SMM, Hendron K, Sloot LH, et al. A soft robotic exosuit improves walking in patients after stroke. Science Translational Medicine. 2017;9(400):eaai9084. doi:10.1126/scitranslmed.aai9084.

12. Turner DL, Ramos-Murguialday A, Birbaumer N, Hoffmann U, Luft A. Neurophysiology of robot-mediated training and therapy: A perspective for future use in clinical populations. Frontiers in Neurology. 2013;4 NOV(November):1–11. doi:10.3389/fneur.2013.00184.

13. Kirtley C, Whittle MW, Jefferson RJ. Influence of walking speed on gait parameters. Journal of Biomedical Engineering. 1985;7:282–288. doi:10.1016/0141-5425(85)90055-X.

14. Riley PO, Croce UD, Kerrigan DC. Propulsive adaptation to changing gait speed. Journal of Biomechanics. 2001;34:197–202. doi:10.1016/S0021-9290(00)00174-3.

15. Monaco V, Rinaldi LA, Macrı G, Micera S. During walking elders increase efforts at proximal joints and keep low kinetics at the ankle. Clinical Biomechanics. 2009;24(6):493–498. doi:10.1016/j.clinbiomech.2009.04.004.

16. Ardestani MM, Ferrigno C, Moazen M, Wimmer MA. From normal to fast walking: Impact of cadence and stride length on lower extremity joint moments. Gait and Posture. 2016;46:118–125. doi:10.1016/j.gaitpost.2016.02.005.

17. Devita P, Copple T, Patterson J, Rider P, Long B, Steinweg K. Modulating step length during walking by young and old adults. In: ASB 2009;.

18. Allet L, Ijzerman H, Meijer K, Willems P, Savelberg H. The influence of stride-length on plantar foot-pressures and joint moments. Gait and Posture. 2011;34:300–306. doi:10.1016/j.gaitpost.2011.05.013.

19. Latt MD, Menz HB, Fung VS, Lord SR. Walking speed, cadence and step length are selected to optimize the stability of head and pelvis accelerations. Exp Brain Res. 2007;184(2):201–209.

20. Hof AL. Scaling gait data to body size. Gait &Posture. 1996;4(3):222–223.

21. Kramer PA, Sarton-Miller I. The energetics of human walking: Is Froude number (Fr) useful for metabolic comparisons? Gait & Posture. 2008;27(2):209–215.

22. Alexander R. The gaits of bipedal and quadrupedal animals. The International Journal of Robotics Research. 1984;.

23. Pataky TC, Vanrenterghem J, Robinson MA. The probability of false positives in zero-dimensional analyses of one-dimensional kinematic, force and EMG trajectories. Journal of Biomechanics. 2016;49(9):1468–1476.

24. Pataky TC, Vanrenterghem J, Robinson MA. Zero-vs. one-dimensional, parametric vs. non-parametric, and confidence interval vs. hypothesis testing procedures in one-dimensional biomechanical trajectory analysis. Journal of Biomechanics. 2015;48(7):1277–1285.

25. Franklin DW, Wolpert DM. Computational mechanisms of sensorimotor control. Neuron. 2011;72(3):425–442. doi:10.1016/j.neuron.2011.10.006.

26. Prochazka A, Gillard D, Bennett DJ. Implications of positive feedback in the control of movement. Journal of Neurophysiology. 1997;77(6):3237–3251.

